# Determinants of stem cell enrichment in healthy tissues and tumors: implications for non-genetic drug resistance

**DOI:** 10.1101/663146

**Authors:** Lora D. Weiss, P. van den Driessche, John S. Lowengrub, Dominik Wodarz, Natalia L. Komarova

**Affiliations:** Department of Mathematics, University of California Irvine, Irvine, CA 92797; Department of Mathematics and Statistics, University of Victoria, Victoria, BC V8W 2Y2, Canada; Department of Ecology and Evolutionary Biology, University of California Irvine, Irvine, CA 92797

## Abstract

Drug resistance is a major challenge for cancer therapy. While resistance mutations are often the focus of investigation, non-genetic resistance mechanisms are also important. One such mechanism is the presence of relatively high fractions of cancer stem cells (CSCs), which have reduced susceptibility to chemotherapy, radiation, and targeted treatments compared to more differentiated cells. The reasons for high CSC fractions (CSC enrichment) are not well understood. Previous experimental and mathematical modeling work identified a particular feedback loop in tumors that can promote CSC enrichment. Here, we use mathematical models of hierarchically structured cell populations to build on this work and to provide a comprehensive analysis of how different feedback regulatory processes that might partially operate in tumors can influence the stem cell fractions during somatic evolution of healthy tissue or during tumor growth. We find that depending on the particular feedback loops that are present, CSC fractions can increase or decrease. We define characteristics of the feedback mechanisms that are required for CSC enrichment to occur, and show how the magnitude of enrichment is determined by parameters. In particular, enrichment requires a reduction in division rates or an increase in death rates with higher population sizes, and the feedback mediators that achieve this can be secreted by either CSCs or by more differentiated cells. The extent of enrichment is determined by the death rate of CSCs, the probability of CSC self-renewal, and by the strength of feedback on cell divisions. Defining these characteristics can guide experimental approaches that aim to screen for and identify feedback mediators that can promote CSC enrichment in specific cancers, which in turn can help understand and overcome the phenomenon of CSC-based therapy resistance.

## Introduction

Healthy tissue is maintained by a small population of tissue stem cells, which self-renew and give rise to transit amplifying cells and differentiated cells that make up the majority of the cell population [1]. Homeostasis is ensured by complex regulatory mechanisms including negative feedback loops [2,3]. Stem cells are thought to be an important target for oncogenic transformation [4], which results from the breaking of homeostatic mechanisms. As the tumor grows, it maintains basic features of the underlying tissue hierarchy [5]. Thus, the bulk of the tumor is made up mostly of more differentiated tumor cells, but maintenance of the tumor is thought to depend on a relatively small fraction of cancer stem cells (CSCs), characterized by the presence of specific markers. The exact proportion of cancer stem cells, however, can vary from tumor to tumor, and, within a tumor, over the time course of growth. CSCs tend to be present in small proportions during earlier phases of tumor growth, and tend to make up larger fractions as the disease progresses, [6,7], a process that can be called CSC enrichment. In addition, therapy can have a significant impact on the CSC fraction [8,9]. Cytotoxic chemotherapeutic agents and radiation kill proliferating DC tumor cells more effectively than quiescent CSCs, and the same has been reported for targeted treatments with small molecule inhibitors [10–12]. Following treatment cessation, re-generation of the differentiated tumor cell population from the activated CSCs can restore the tumor composition with a majority of differentiated cells, although there is indication that the fraction of CSCs can in some cases remain permanently elevated post-compared to pre-treatment [13].

Since CSC are less susceptible to therapy than more differentiated cells, the fraction of CSCs in the tumor can be an important determinant of the response to cancer therapy. In particular, a relatively high CSC fraction at the time of treatment can be a cause of non-genetic drug resistance, where a loss of response is observed despite the absence of any known drug-resistant mutants. Thus, in bladder cancer, it has been argued that CSC enrichment might account for the progressive loss of treatment response with repeated cycles of chemotherapy [8]. In chronic myeloid leukemia, high fractions of undifferentiated cells, especially during advanced stages of progression, might contribute to the phenomenon of primary resistance against tyrosine kinase inhibitors [14]. In mouse mammary tumors, CSC enrichment has been shown to contribute to chemotherapy resistance [9]. Hence, understanding the conditions under which stem cell enrichment, and hence high fractions of CSCs, are achieved is paramount to finding strategies to overcome this form of resistance.

The factors that determine the fraction of stem cells in healthy tissues and tumors are not well understood, and neither are the factors that contribute to CSC enrichment during tumor growth, or to sustained CSC enrichment following treatment cycles. Recent experimental and mathematical work implicates the presence of feedback regulatory loops in the process of stem cell enrichment in tumors. For example, a PGE2-mediated wound-healing type response has been documented in bladder cancer that can result in CSC repopulation during chemotherapy [8,15]. According to this mechanism, chemotherapy-induced death of differentiated cells results in the release of positive feedback signals that drive CSCs into a state of proliferation. Mathematical modeling work has shown, however, that CSC enrichment cannot be sustained after treatment cessation by the wound-healing response alone, but might further require the presence of negative feedback loops within the tumor [13]. It was shown mathematically that the presence of negative feedback from differentiated cells onto the rate of CSC division promotes sustained CSC enrichment post therapy, while this is not observed in the absence of negative feedback [13]. The model suggests that enriched CSC populations are only maintained post-therapy if feedback mechanisms also act during untreated tumor growth.

The above-mentioned positive feedback wound-healing mechanism during chemotherapy is to our knowledge the only regulatory feedback loop that has been identified experimentally in tumors. Analysis of untreated tumor growth dynamics, however, has suggested that negative feedback processes might play an important role as well [16,17]. Experimentally, negative feedback has been reported, e.g., in the olfactory epithelium, where GDF11 and Activin βB negatively regulate cell division rates in progenitor and stem cells [2,18]. Another prominent example of a feedback implicated in carcinogenesis is the dual tumor-promoter/suppressor role of TGFβ signaling [19], whereby in early-stage tumors, it promotes cell cycle arrest and apoptosis, and at advanced stages, the TGFβ pathway in fact enhances tumor progression by promoting cancer cell motility, invasions, and “stemness” [20–22]. Therefore, further investigation is warranted into the role of feedback loops for stem cell enrichment, which is the subject of our study.

In our previous work, we only focused on one particular negative feedback loop and showed that it can contribute to stem cell enrichment, i.e. negative feedback from differentiated cells onto the rate of stem cell division. There are, however, a large number of possible negative and positive feedback loops that can potentially influence the proportions of stem cells and differentiated cells in healthy and tumor tissue. These include negative or positive control of rate functions (such as division rate and death rate) and differentiation probability by stem cells, differentiated cells, or both, see [23–25]. The question thus arises, which of these feedback mechanisms can drive stem cell enrichment, and which cannot. Finding an answer to this question is the subject of our investigation and might help to find treatment methods that can sensitize tumors to chemotherapies and targeted therapies.

We focus on two basic scenarios. First, we assume that partial breakage of feedback regulation results in temporary cell growth towards a new equilibrium, characterized by an overall larger number of cells. This could correspond to a single step in step-wise tumor progression. We investigate the conditions required for the stem cell fraction to be larger at the new compared to the old equilibrium. In particular we study how remaining feedback loops determine the stem cell fraction. Second, we consider unbounded tumor growth and investigate how different feedback mechanisms that remain in a growing tumor cell population can determine whether or not CSC enrichment occurs during growth. Much of this work is done using ordinary differential equations. In the context of unbounded tumor growth, the effect of spatial growth patterns on stem cell enrichment is explored using a stochastic agent-based model and an analytical approximation.

### The basic mathematical modeling approach

An ordinary differential equation model has been used to describe tissue hierarchy dynamics in a healthy tissue [2,26], and the models presented here build on these approaches. While cell lineages consist of stem cells, transit amplifying cells, and terminally differentiated cells, our models make a simplification and take into account only stem cells (which encompass all the proliferating cells) and differentiated cells. Denoting stem cells (SC) by x and differentiated cells (DC) by y, the model is given by:

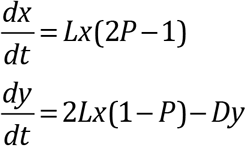

Stem cells divide with a rate L. With a probability P, the division results in two daughter stem cells (self-renewal), and with a probability (1-P), the division results in two daughter differentiated cells (differentiating division). Differentiated cells are assumed to die with a rate D. This model captures a probabilistic model of tissue control, which occurs on the population level if about half of the symmetric divisions result in two daughter stem cells and the other half result in two daughter differentiated cells. In addition to symmetric divisions, asymmetric divisions may play a role in tissue renewal. With asymmetric cell division, a stem cell gives rise to one stem cell and one differentiated cell, thus maintaining a constant population of stem cells. In the current model, even though such divisions do not appear explicitly, it is possible to show that they are included implicitly, see Section 1.2 of the Supplement. It is proven that a model that considers both symmetric and asymmetric divisions is mathematically identical to the one studied here.

The basic model shown above contains no feedback loops. Hence, the rates L and D and probability P are constants that are independent of x or y. This system is only characterized by a neutrally stable family of nontrivial equilibria if P=0.5. If P>0.5, infinite growth is observed. If P<0.5, the cell population goes extinct.

It has been shown that introduction of negative feedback loops can result in more realistic behavior, where a stable equilibrium is attained for P>0.5 [2]. This was shown in the context of two specific feedback loops, and subsequently generalized to comprehensively list all possible (positive and negative) feedback loops compatible with stability [23–25]. Here, we also use a general model to assume different kinds of feedback on the rate of cell division, L, the rate of cell death, D, and the probability of self-renewal, P. We also add the possibility that stem cells die with a rate δ (which can also be subject to feedback). In the context of our model, feedback is equivalent to a dependence of rates and probabilities on the population sizes, x and/or y. Hence, the model is given by the following ODEs:

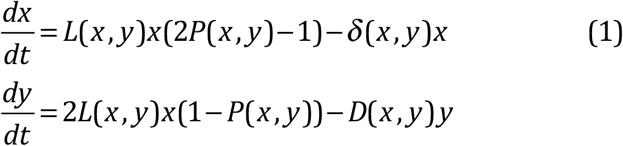

The division rate L, the death rates, D and δ, and the probability of self-renewal, P, are now functions of either the number of stem cells, x, or the number of differentiated cells, y, or both.

Evolution can result in the generation of mutant cell populations that are characterized by a higher self renewal probability, given by P_2_. Hence, we now have two stem and differentiated cell populations denoted by subscripts 1 and 2 for wild type and mutant types, respectively. The equations are thus given by

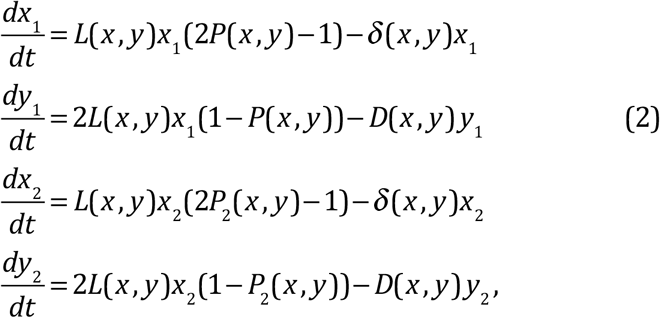

where *x*=*x*_1_+*x*_2_ and *y*=*y*_1_+*y*_2_. The two cell populations are in competition with each other, mediated by the feedback factors that are shared between the two populations. Table 1 summarizes all the variables used in this paper (both in this section and in the later sections).

### Cell growth towards a new equilibrium

The first important scenario happens when the mutant cell population gains a selective advantage, outcompetes the original, healthy cell population, and grows towards a new and higher equilibrium level. This is achieved by assuming that P_2_>P, along with other conditions on the rate functions that are specified in Section 1 of the Supplement. We examine the conditions under which the stem cell fraction at the new equilibrium is increased compared to that at the original equilibrium. In terms of the model notation, we are interested in the quantity v(x,y) = x/ y, i.e. the ratio of stem to differentiated cells. We investigate how different combinations of feedback loops that remain in the mutant cell population impact stem cell enrichment.

**Table 1.**
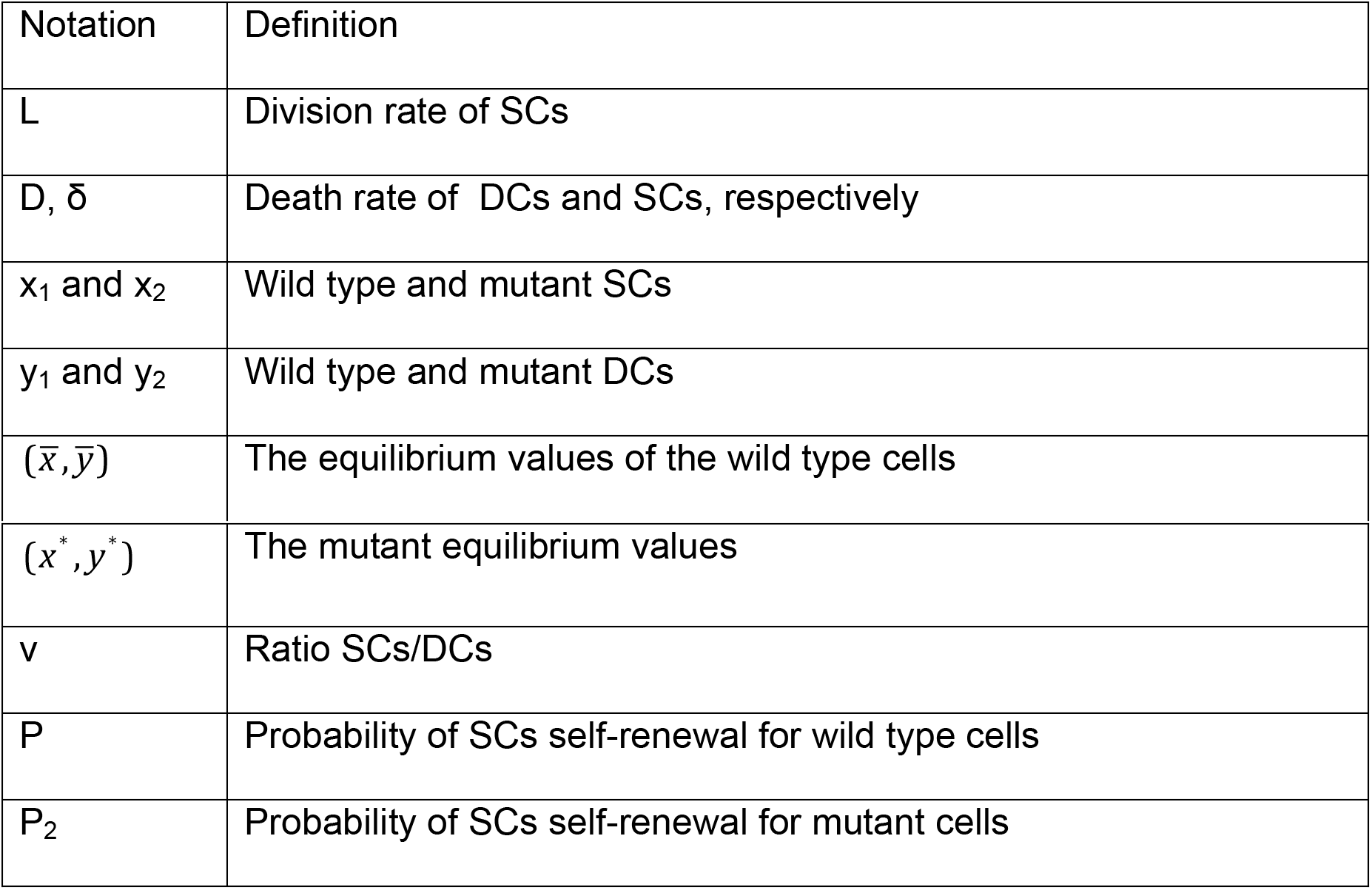
Mathematical symbols and the definitions of variables.

| Notation | Definition |
| --- | --- |
| $L$ | Division rate of SCs |
| $D, \delta$ | Death rate of DCs and SCs, respectively |
| $x_1$ and $x_2$ | Wild type and mutant SCs |
| $y_1$ and $y_2$ | Wild type and mutant DCs |
| $(\bar{x}, \bar{y})$ | The equilibrium values of the wild type cells |
| $(x^*, y^*)$ | The mutant equilibrium values |
| $v$ | Ratio SCs/DCs |
| $P$ | Probability of SCs self-renewal for wild type cells |
| $P_2$ | Probability of SCs self-renewal for mutant cells |

In the following we assume that the division and death rates in the above models are monotonic functions of the number of stem and/or differentiated cells. We further assume that in the absence of mutants, the system is at equilibrium, characterized by the pair 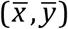, which satisfies:

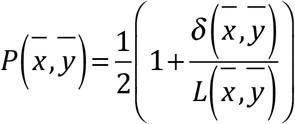

At this equilibrium, the fraction of stem to differentiated cells is given by

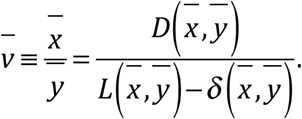

We assume that the mutant cell population can invade from low numbers and displace the original cell population. This occurs if P_2_(x,y) > P(x,y) (this is a sufficient condition). Assuming that a new equilibrium is reached, it is characterized by 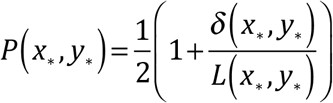
and the new fraction of stem cells is given by

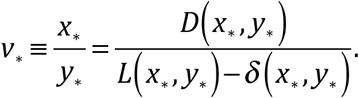

We examine under what conditions 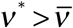, i.e. when the stem cell ratio at the newly obtained mutant equilibrium exceeds that at the original equilibrium.

We observe two qualitatively distinct outcomes, see Sections 2 and 3 of the Supplement for the detailed analysis. (i) The fraction of stem cells increases compared to the previous equilibrium state. We call this stem cell enrichment. This occurs if the ratios L/D and/or L/ δ are decreasing functions of the cell population. This happens when larger cell population sizes result in negative feedback on cell expansion parameters and/or positive feedback on death. For example, this can occur if the per cell division rate decreases and/or the death rate increases with population size. (ii) The fraction of stem cells decreases compared to the previous equilibrium state. We call this stem cell depletion. This occurs if the ratios L/D or L/ δ are increasing functions of the cell population. In this scenario, larger populations promote cell expansion kinetics, e.g. by decreasing the death rate of cells or increasing their division rate. This would correspond to a positive feedback loop on cell division and/or a negative feedback loop on death rate. We note that mutant emergence generally changes the stem cell fraction, unless the division and death rates (L, D, and δ) are constant and hence not affected by feedback.

We will illustrate these points by using some specific examples. The first example is of SC enrichment. Consider a system where δ=0, the self renewal probabilities P and P_2_ are given by decreasing functions of y (see solid lines in Fig 1(a)), and the division rate L is also a function of y, solid line in Fig 1(b). The cell dynamics and control loops corresponding to this system are schematically shown in Fig 2(a). In this and other such diagrams, blue arrows correspond to cellular processes (characterized by kinetic rates, such as L and D), and cell fate decisions, which are probabilities (P or P_2_). For example, the horizontal arrow connecting a circle marked “SC” with the circle marked “?” denotes a SC division. The question mark reminds the reader that a cell fate decision needs to be made, whether to self-renew (a blue arrow going back to SC, marked with probability (1-P)), or to differentiate (a horizontal arrow toward a circle marked with DC, probability P). This notation allows us to show precisely which populations control which processes. In Fig 2(a), the red negative arrows originating in the DC circle represent a negative dependence of both functions L and P on y (the number of DCs).

**Fig 1:**
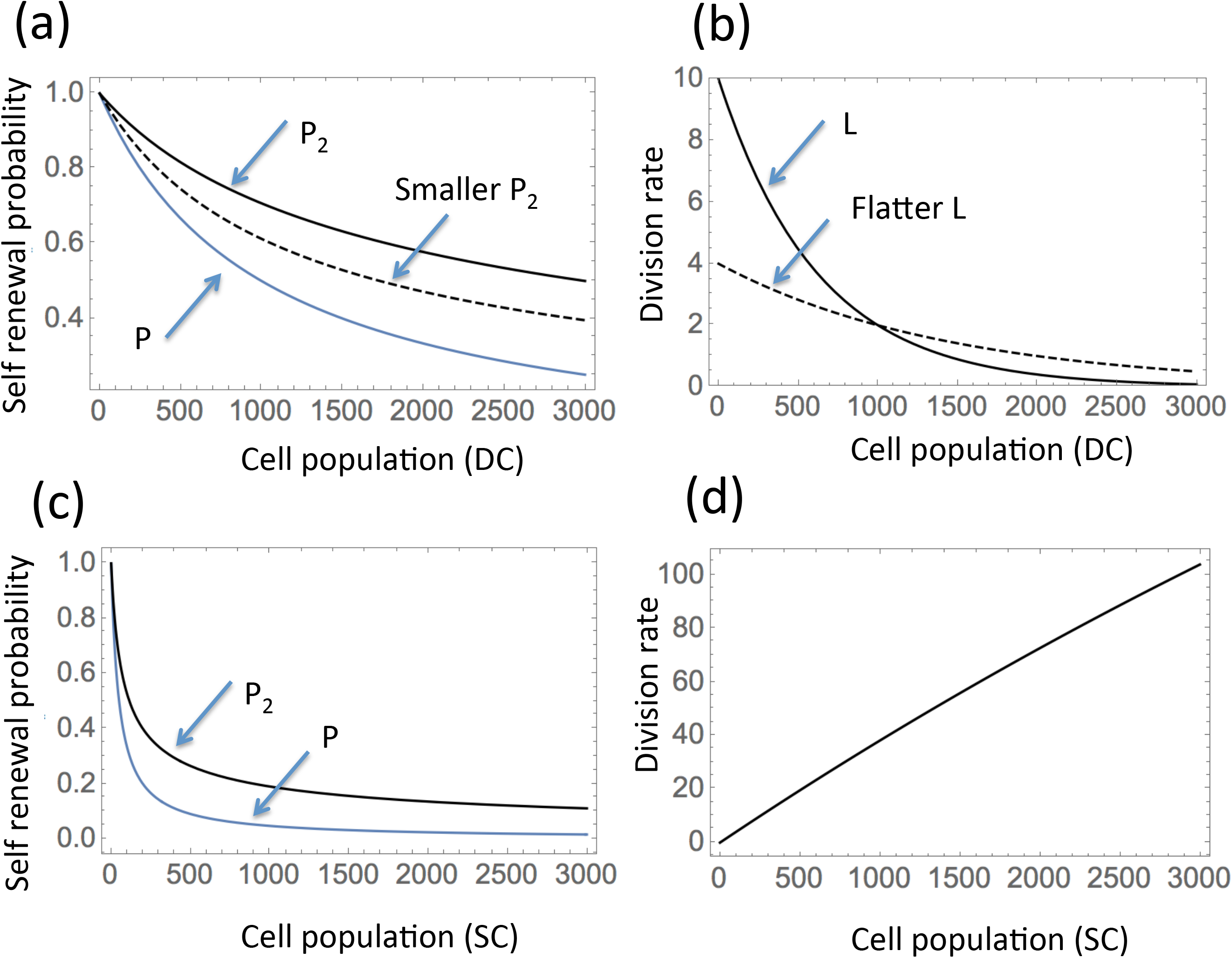
Examples of functional dependencies leading to SC enrichment (a,b) and depletion (c,d) at a new equilibrium. (a) Self renewal probability functions, P(y)=(1+0.001 y)^−1^ and P_2_(y)=(1+0.001 y)^1/2^ P(y) (the solid lines). An example of a smaller self renewal probability of mutants is given by P_2_(y)=(1+0.005y)^1/2^ P(y) (the dashed line). (b) An example of a negatively controlled division rate is given by L(y)=2×5^1-0.001y^ (solid line). A flatter division rate is given by L(y)=2^2-0.001y^ (dashed line). (c) Self renewal probability functions, P(x)=(1+0.02 x)^−1^ and P_2_(x)=(1+0.015 x)^1/2^ P(x). (d) An example of a positively controlled division rate is given by L(x)=401(1-e^−0.0001x^).

**Fig 2:**
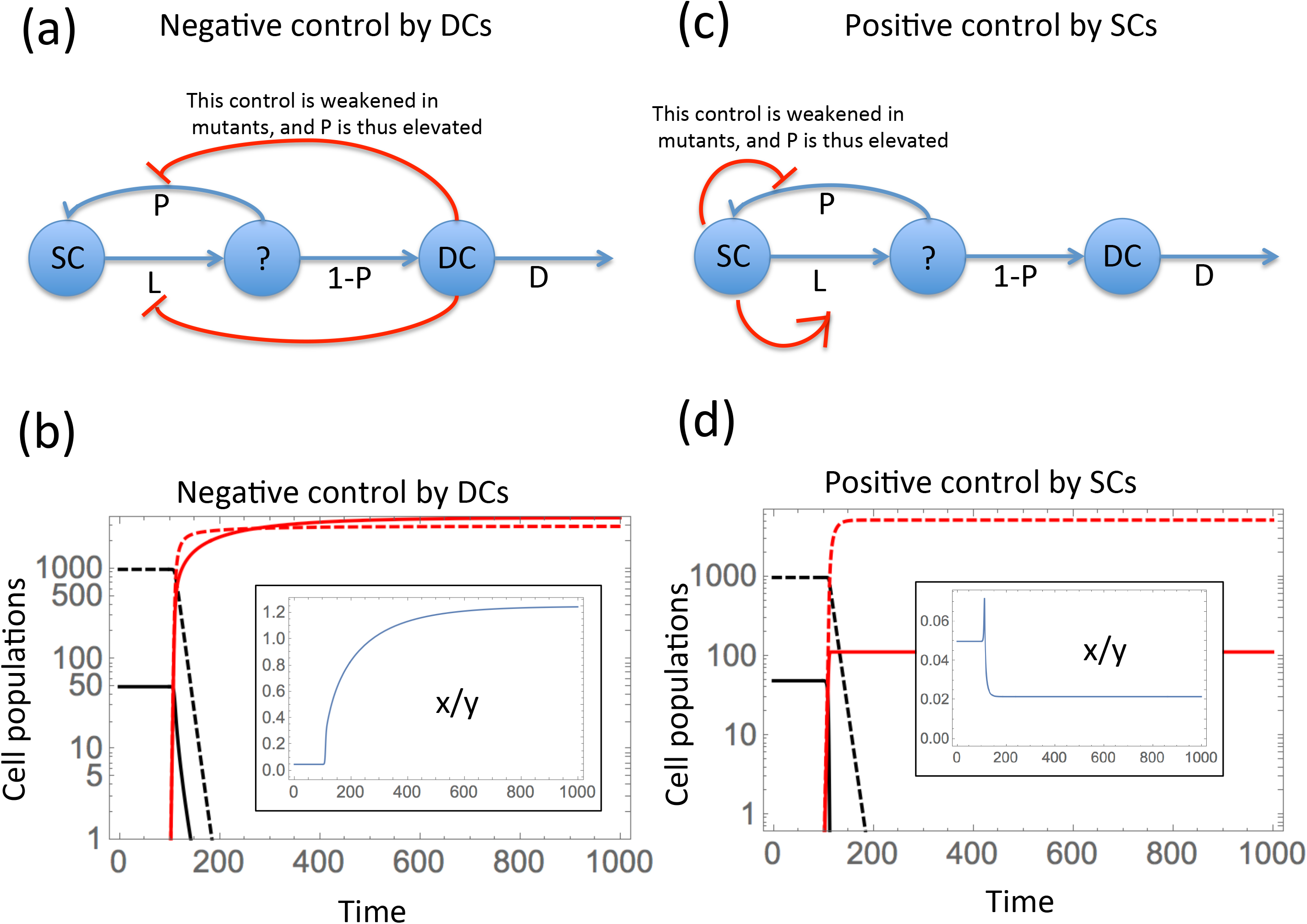
Dynamics of enrichment when a new equilibrium is reached by mutants. The left panels (a, b) illustrate SC enrichment, and the right panels (c,d) SC depletion. In the top panels, the dynamics and the control loops are presented schematically. SCs (the leftmost circle) divide at rate L, such that the decision of whether to self renew or to differentiate (denoted by a question mark) is governed by probability P. Control loops are depicted by red arrows (positive or negative) directed from the population mediating the control to the rate/probability that is being controlled. In the bottom panels, the time series are shown, where functions for wild type cells, x_1_(t) and y_1_(t), are plotted in black, and functions for mutant cells, x_2_(t) and y_2_(t), are plotted in red. SCs are depicted by solid and DCs by dashed lines. Initially, the system is at the original equilibrium, 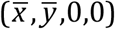. At t=100, mutant stem cells are introduced at a low level, resulting in the extinction of wild type cells, and convergence to a new equilibrium, (0,0,x_*_, y_*_). In the insets, the SC fraction, x/y, is depicted as a function of time. (a,b) Negative control by DCs, resulting in SC enrichment: the functions L, P, and P_2_ are given by the solid lines in Fig 1(a,b). (c,d) Positive control by SCs, resulting in SC depletion: the functions L, P, and P_2_ are specified in Fig 1(c,d).

In this first example, SC enrichment is predicted to occur. Fig 2(b) shows the cell dynamics, once a mutant is introduced at 100 time units. While the wild type population goes extinct, the mutants rise to a new equilibrium characterized by a significantly higher ratio x/y compared to the original equilibrium.

The second example is SC depletion. It is given by a system with rate functions given by Fig 1(c,d), with δ=0 and the division rate positively controlled by the SC population. The controls are schematically shown in Fig 2(c). The resulting dynamics are presented in Fig 2(d), where the proportion of SCs at the new, mutant equilibrium is smaller than the original proportion. Note however that the dependence of x/y on time is non-monotonic and a temporary phase of SC enrichment is experienced before the ratio x/y lowers to its long-term level.

These two examples illustrated in Fig 2 show that the proportion of SCs can either increase (SC enrichment) or decrease (SC depletion), once mutants with altered (increased) self-renewal probability take over the cell population. We refer to Section 2 of the Supplement for the detailed analysis of the scenario where all the rates and functions are control either by SCs or by DCs, but not both. Section 3 of the Supplement extends this to the more general case where the rate functions depend both on x and y.

Next, we examine the factors that affect the magnitude of the change. In Fig 3 we take the basic model of Fig 2(a) and modify it in three different ways, each of which leads to a decrease in the amount of SC enrichment, compared to the situation in Fig 2(a). First, we consider the death rate of the stem cells, δ. The amount of enrichment is smaller for nonzero stem cell death rates, compared to the case of δ=0. In Fig 3(b) we can see that in the presence of SC death, the increase in the enrichment parameter, x/y, is more modest than that of Fig 2(b), 0.4 compared to 1.2.

**Fig 3.**
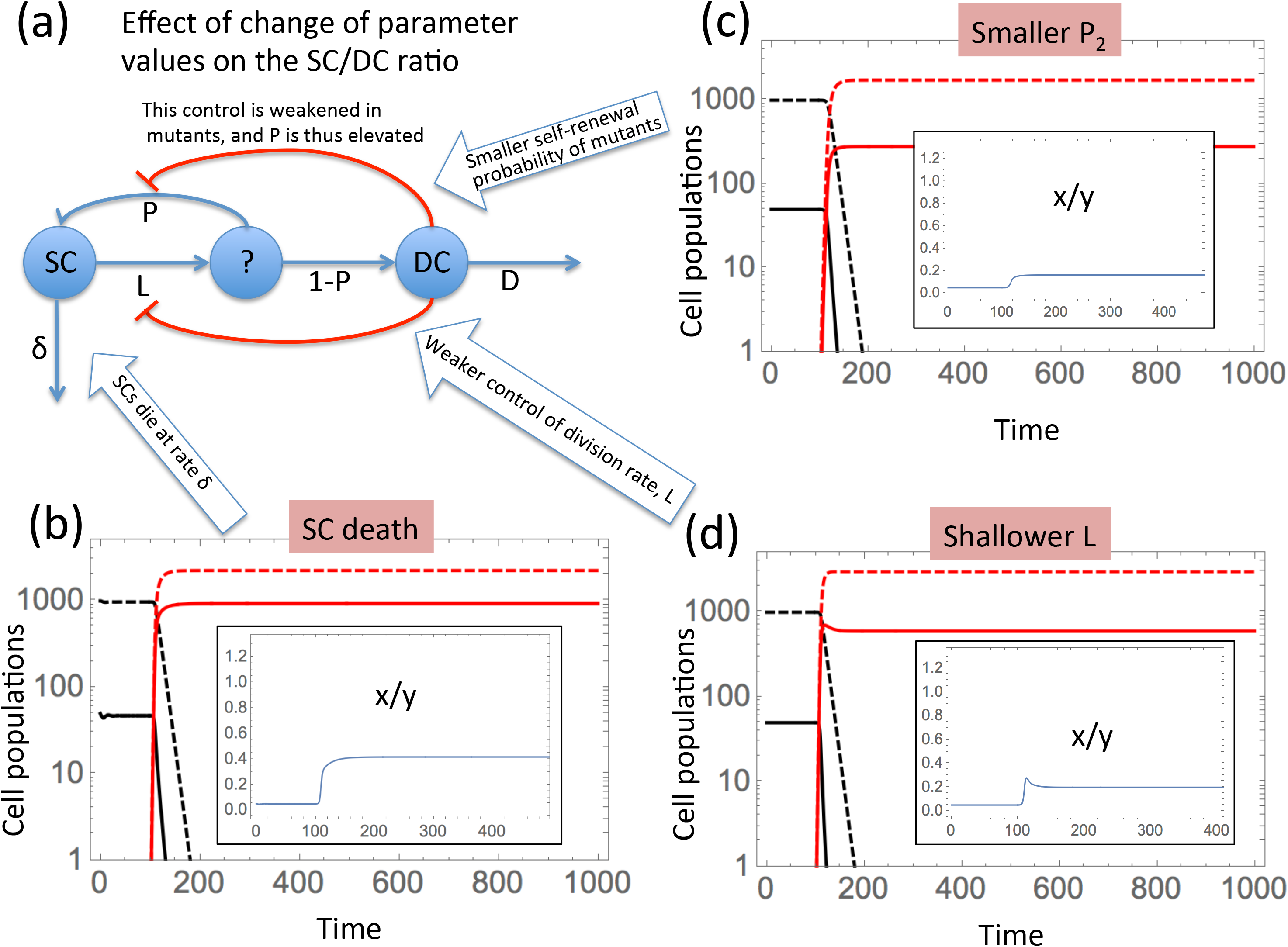
Parameter dependence of the SC enrichment magnitude. (a) The basic model, whose 3 aspects are modified. (b) SC death: same parameters as in Fig 2(a), except δ=0.03. (c) Smaller P_2_: same parameters as in Fig 2(b) except P_2_ is given by the dashed line in Fig 1(a). (d) Shallower L: same parameters as in Fig 2(b), except the division rate L is given by the dashed line in Fig 1(b). We observe that the SC enrichment is (b-d) is less pronounced compared to Fig 2(b).

The second modification is a smaller self-renewal probability, P_2_, of the mutant cell population, Fig. 3(c). The self-renewal probability P_2_ of the mutant cells is that given by the dashed line in Fig 1(a). Again, this results in a more modest increase in the stem cell fraction compared to Fig 2(b), 0.1 vs 1.2.

The third modification is a less pronounced feedback on the division rate. The result is that the SC enrichment becomes smaller. In Fig 3(d) we use a flatter division rate function, L, than that in Fig 2(a) (compare the dashed line in Fig 1(b) with the solid line). This results in a smaller SC enrichment, as shown by a smaller increase of x/y in Fig 3(d) (0.2 compared to 1.2 of Fig 2(b)). A detailed analysis of all these scenarios is presented in Section 2 of the Supplement.

### Stem cell enrichment in non-equilibrium situations

Next, we study the scenarios where the mutant population grows from low numbers and does not reach a new equilibrium, but instead, continues to grow indefinitely. This would correspond to tumor growth. The definition of enrichment in this context and the relevant methodology will be somewhat different in this case. We will consider the mutant population alone, and study the growth of x_2_ and y_2_, in order to find the dynamics of the quantity ν=x_2_/y_2_. We will say that stem cell enrichment occurs if the quantity ν increases during population growth, either infinitely, or temporarily. For simplicity we will assume that the cancer stem cell (CSC) population does not die (δ=0). We further assume that the probability of self-renewal of mutants, P_2_, is a monotonic function of the population size (stem cells or differentiated cells) that satisfies P_2_ > 1/2, and that in the limit of large populations, it approaches a limiting value, 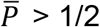. When we consider the mutant dynamics, we will drop the subscript 2, and simply study the equations:

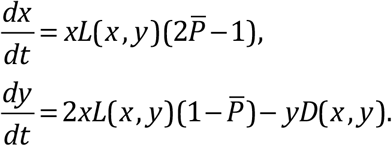

If we assume that both D and L depend on a single population, that is, L=L(x), D=D(x), or L=L(y),D=D(y), the following approximations can be derived (see Section 4 of the Supplement). As the cell population expands, the DC population behaves as

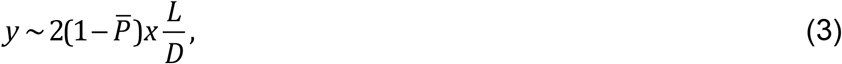

and ratio of stem to differentiated cells, v, in the limit of large times is given by:

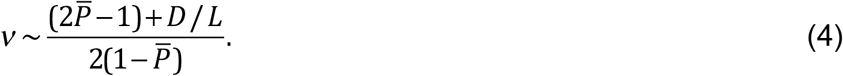

Equation (4) states that the long-term dynamics of the SC fraction, v, is defined by the behavior of the ratio D/L, the per cell death rate of the DCs and per cell division rate of SCs. If either or both of these quantities are controlled by the population size, the ratio will change as the tumor grows. If the behavior of the death rate and division rates are known, formula (4) predicts the dynamics of the SC fraction in tumor growth. Three scenarios are possible.

#### 1) Unlimited SC enrichment

One relevant scenario occurs if the ratio D/L grows indefinitely as the population size increases; this corresponds to either a negative feedback on L (such that L approaches zero as population size increases), and/or a positive feedback on D. If an increase in cell population sizes results in an unbounded increase in D/L, then the stem cell fraction, v, will continuously rise towards infinity. Therefore, as the tumor cell population grows, the fraction of SCs increases, and at very large tumor sizes, the tumor is practically only made up of CSCs. This scenario is illustrated in figures 4(a,b) and 5(a,b), where negative feedback on L results in an increase in the ratio D/L, mediated by differentiated cells (panels (a)) and stem cells (panels (b)). Figs 4(a), 5(a) explores negative feedback on L by DCs. It depicts a system similar to that of Fig 2(a), except the self-renewal probability of the mutants is a constant, resulting in an unlimited growth of mutants. The fraction x/y in this case experiences unlimited growth, as shown by the inset in panel 5(a). In Figs 4(b) and 5(b), we replace control of L by DCs with control by SCs; again, an unlimited growth of x/y is observed, see the inset in panel 5(b).

**Fig 4.**
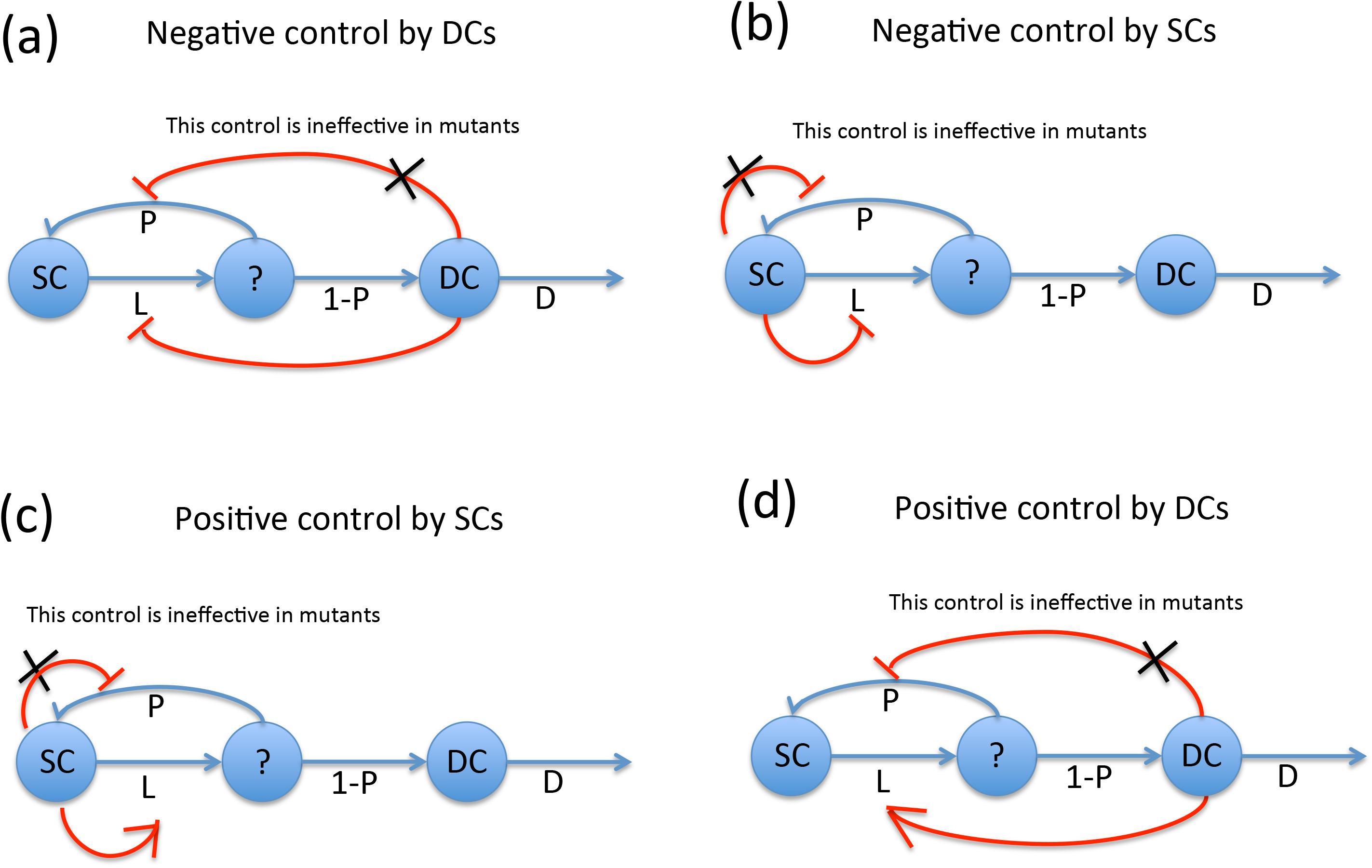
Four types of control used in Fig 5 to study unbounded growth. (a) Negative control by DCs; (b) Negative control by SCs; (c) Positive control by DCs; Positive control by SCs; (d) Positive control by SCs. Unlike in Fig 2(a,c), the mutant self-renewal probability is assumed to be constant (and thus the control loop is completely severed in mutants), leading to unlimited growth. The 2 top panels correspond to Fig 5(a,b); the 2 bottom panels correspond to Fig 6(a,b).

**Fig 5.**
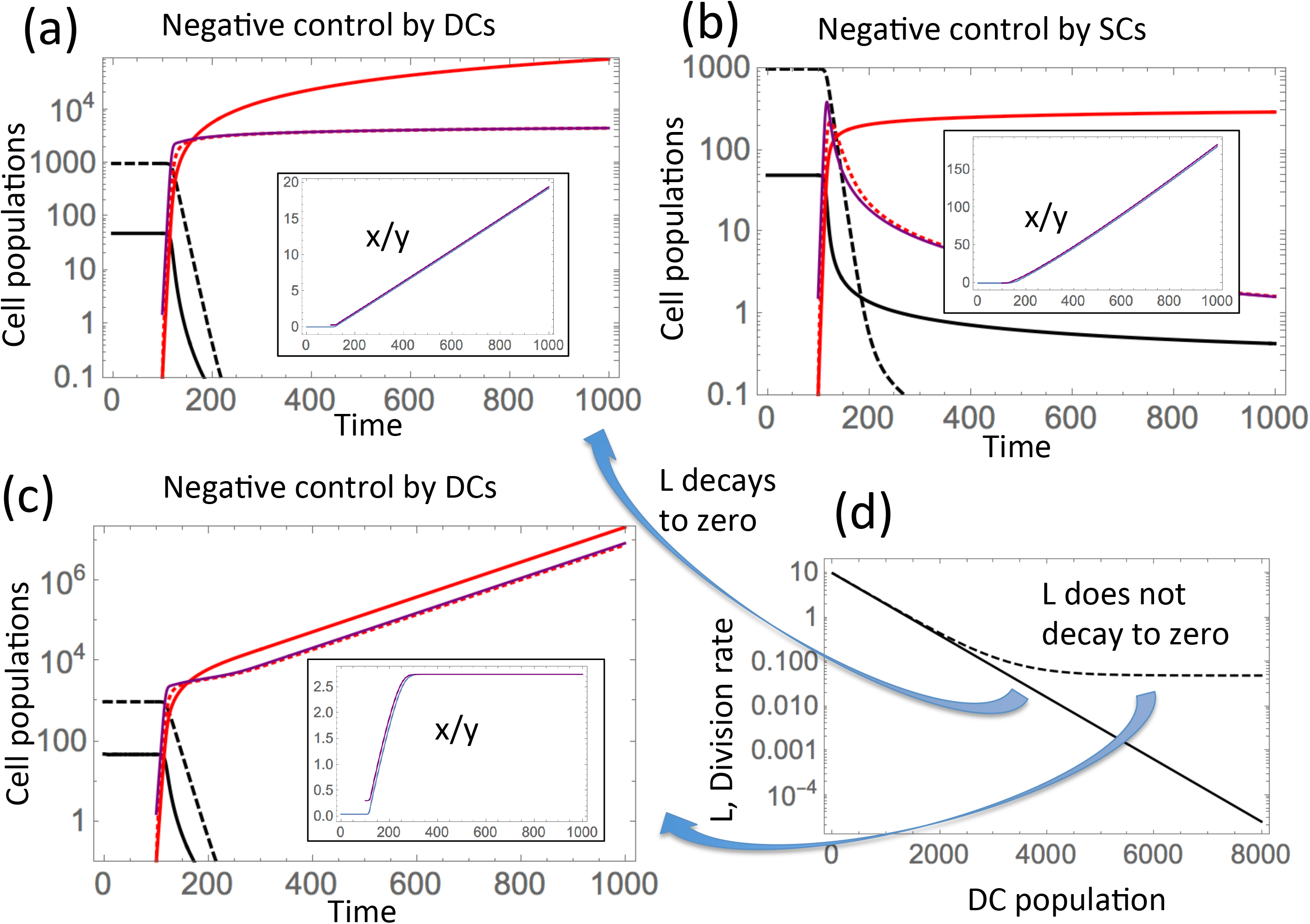
Dynamics of unbounded growth under negative control on SC divisions. Notations are as in Fig 2(b,d). The purple line in each panel plots the approximation for y (formula (3)), and in the insets the approximation for x/y (formula (4)). (a) Negative control by DCs: parameters are as in Fig 2(b) except 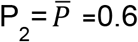. (b) Negative control by SCs: similar to (a), except the rate functions depend on SCs: L(x)=2*5^(1-x/50)^, P(x)=(1+0.02x)^−1^, and 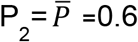. In both panes, unlimited SC enrichment is observed. (c) Saturated SC enrichment: same as (a), except the division rate decreases to a constant: L(y)=0.05+2*5^1-0.001y^. (d) The two division rates are plotted on a logarithmic scale as functions of DC populations. The solid line depicts the function used in panel (a) and the dashed line the function used in panel (c).

**Fig 6.**
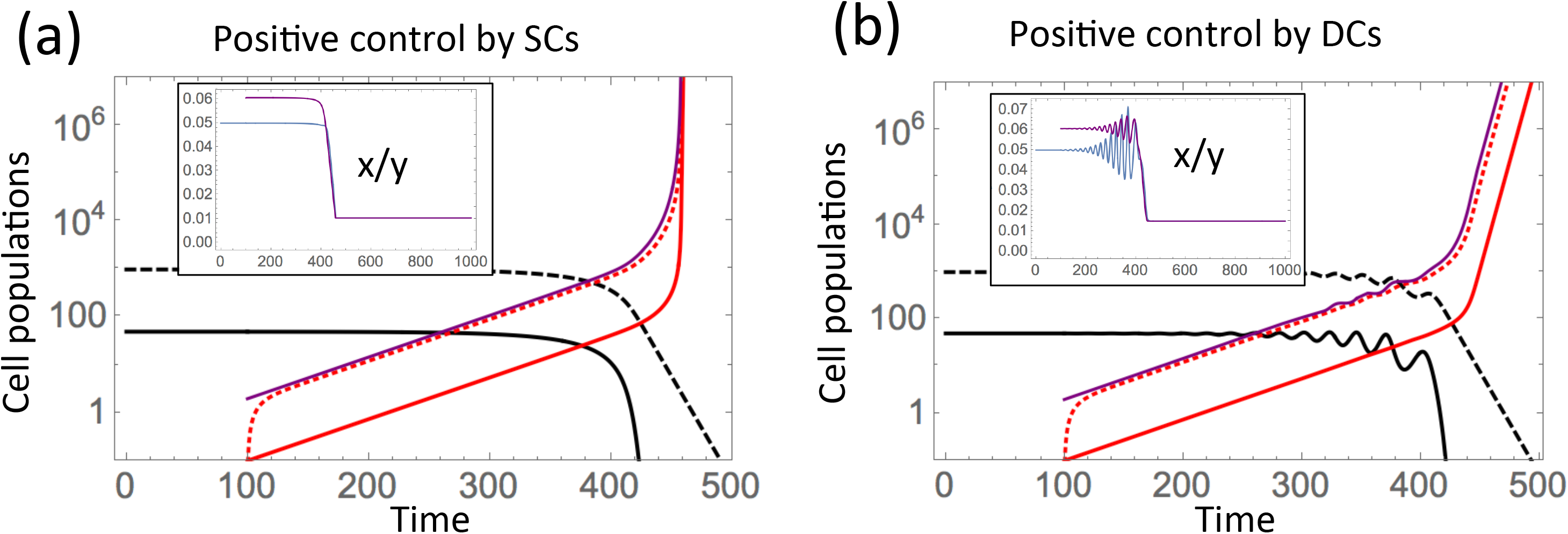
Dynamics of unbounded growth under positive control on SC divisions. Notations are as in Fig 2(b,d). The purple line in each panel plots the approximation for y (formula (3)), and in the insets the approximation for x/y (formula (4)). (a) Positive control by SCs: corresponds to parameters of Fig 2(d), except 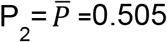. (b) Positive control by DCs: similar to (a), except the rate functions depend on DCs: L(y)=21(1-e^−0.0001y^), P(y)=(1+0.001y)^−1^, 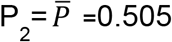.

#### 2) Saturated SC enrichment

The second type of behavior is observed if an increase in the number of cells results in a saturated growth in D and/or a decay of L to a nonzero level, such that the ratio D/L is a growing and saturating function as the cell population increases. In this case, the ratio v will increase and reach a constant level, as the tumor size grows beyond a given size. This scenario is illustrated in figure 5(c,d), where populations x and y grow indefinitely, but the ratio x/y first increases and then reaches a constant level (see the inset). The only difference in simulations of panel 5(c) compared to those of panel 5(a) is the fact that the division rate, L, no longer decreases to zero, but reaches a small but nonzero constant level, as the cell population grows. Panel 5(d) depicts the two functions, L(y), that are used in the simulation (the solid line for panel 5(a) and the saturating, dashed line for panel 5(c)).

#### 3) Saturated SC depletion

The third and final scenario that is possible corresponds to the case where function D/L decreases as the population size increases. This can happen if the division rate is positively controlled and/or when the death rate is negatively controlled by the cell populations. This type of dynamics is illustrated in figures Fig 4(c, d) and 6(a,b). In these simulations, positive feedback on L was mediated either by SCs (figures 4(c) and 6(a)) or by differentiated cells (figures 4(d) and 6(b)). In this case, a temporary reduction in the ratio ν can be observed during tumor growth, until ν converges to a constant. In other words, we observe a reduction in the CSC fraction over time, until it reaches a limiting value. Note that the SC fraction can never decrease to zero, the minimum fraction is given by 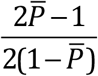.

To summarize, the following three types of SC fraction dynamics are predicted: (1) If L is subject to negative feedback and decays to zero and/or D is subject to positive feedback and increases without bound, then we have unlimited SC enrichment. (2) If L is subject to negative feedback but never decreases to zero and/or D is subject to positive feedback and increases within bounds, then we have limited SC enrichment, where the SC fraction increases and then remains constant. (3) If L is subject to positive feedback and/or D is subject to negative feedback, then we have SC depletion, and the SC fraction decreases to a nonzero level.

Note that if L and D are subject to the same type of feedback, then the resulting behavior would be determined by the overall behavior of L/D. For example, if L and D are both subject to positive feedback and increase within bounds, but the feedback on L is greater than the feedback on D such that L/D is increasing with the population size, then this would correspond to type (3) and result in depletion. However, if the feedback on D is (effectively) greater such that L/D is decreasing and bounded above 0, then this would correspond to type (2) and result in limited SC enrichment.

### SC enrichment in spatially structured populations

According to the above results, stem cell enrichment during growth requires certain feedback mechanism to be present in the tumor cell population, such that the ratio L/D is reduced as the population size is increased. Typically such feedback can occur through signaling factors that are secreted from stem or differentiated cells. Another way to achieve a similar result can be spatially restricted reproduction of cells. In such scenarios, cells experience range expansion in two dimensions, or grow as expanding sphere-like structures in 3D. Inside the expanding population, divisions must be balanced with deaths because free space is limited. In a way this works similarly to control loops affecting division and/or death rates, which were discussed earlier in the paper; in the case of spatially restricted growth, control is essentially competition for space, which leads to slower divisions/ higher death as the density increases.

To explore this, we first considered a two-dimensional stochastic agent-based model that describes spatially restricted cell growth. The model assumes a 2-dimensional grid consisting of nxn spots. A spot can either be empty, contain a stem cell, or contain a differentiated cell. At each time step, the grid is sampled N times, where N is the number of cells currently present in the grid. If the sampled spot contains a stem cell, it divides with a probability L_0_, and dies with a probability δ_0_. If the division event is chosen, one of the eight nearest neighboring spots is randomly picked as a target for one of the daughter cells. If the chosen spot is already filled, the division event is aborted, otherwise it proceeds. If division proceeds, both daughter cells will be stem cells with a probability P_0_ (self-renewal). With probability 1-P_0_, both daughter cells will be differentiated cells. If the sampled spot contains a differentiated cell, death occurs with a probability D_0_. No explicit feedback processes were included in the model. The simulation was started with 9 stem cells (and no differentiated cells). Assuming that stem cells do not die, the resulting average growth curve is shown in Figure 7(a). While initially, the differentiated cells grow to be more abundant than the stem cells, the stem cell population enriches over time and eventually becomes dominant as the cell population grows. Figure 7(b) shows the same kind of simulation, but assuming that stem cells die with a rate that is smaller than the death rate of differentiated cells. Consistent with the results obtained for explicit feedback mechanisms, we find that the degree of stem cell enrichment is reduced in the presence of stem cell death (higher rates of stem cell death lead to less enrichment).

**Figure 7.**
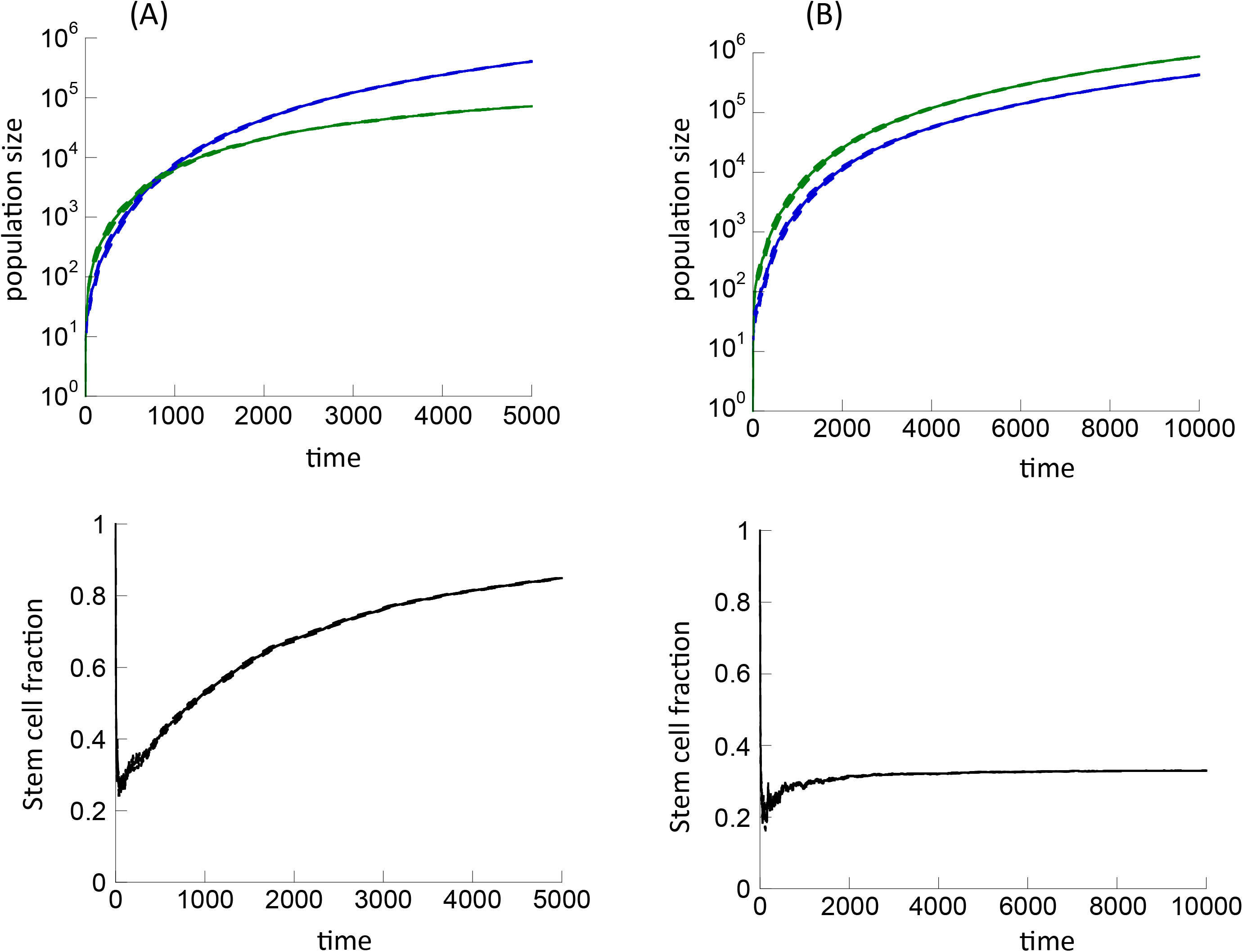
Effect of spatially restricted cell division on CSC enrichment, according to the 2-dimensional implementation of the agent-based model. (A) Dynamics in the absence of stem cell death, δ_0_=0. In the top panel, stem cells are shown in blue, differentiated cells in green. The bottom panel shows the stem cell fraction over time. (B) Same, but for a non-zero stem cell death rate, i.e. δ_0_=0.005. Other parameters were chosen as follows: L_0_=0.95, P_0_=0.6, D_0_=0.01, nxn=1500^2^. 8 simulations were run, and the mean is shown as a solid line while the mean ± standard error are shown by dashed lines.

A simple mean-field model that takes account of the space limitations inside an expanding population can explain these results (see Section 5 of the Supplement). Indeed, the system in the interim of the expanding globe reaches a dynamic equilibrium state where the density of SCs and DCs is dictated by the balance of division and death rates. Figure 8 shows theoretical predictions for the densities of SCs and the DCs in the colony’s interim; the solid lines depict theoretical predictions and points correspond to numerical simulations of the agent based model. The figure also provides typical images for two parameter combinations; blue dots represent SCs and yellow dots DCs. As expected, under higher DC death the proportion of DCs decreases.

**Figure 8.**
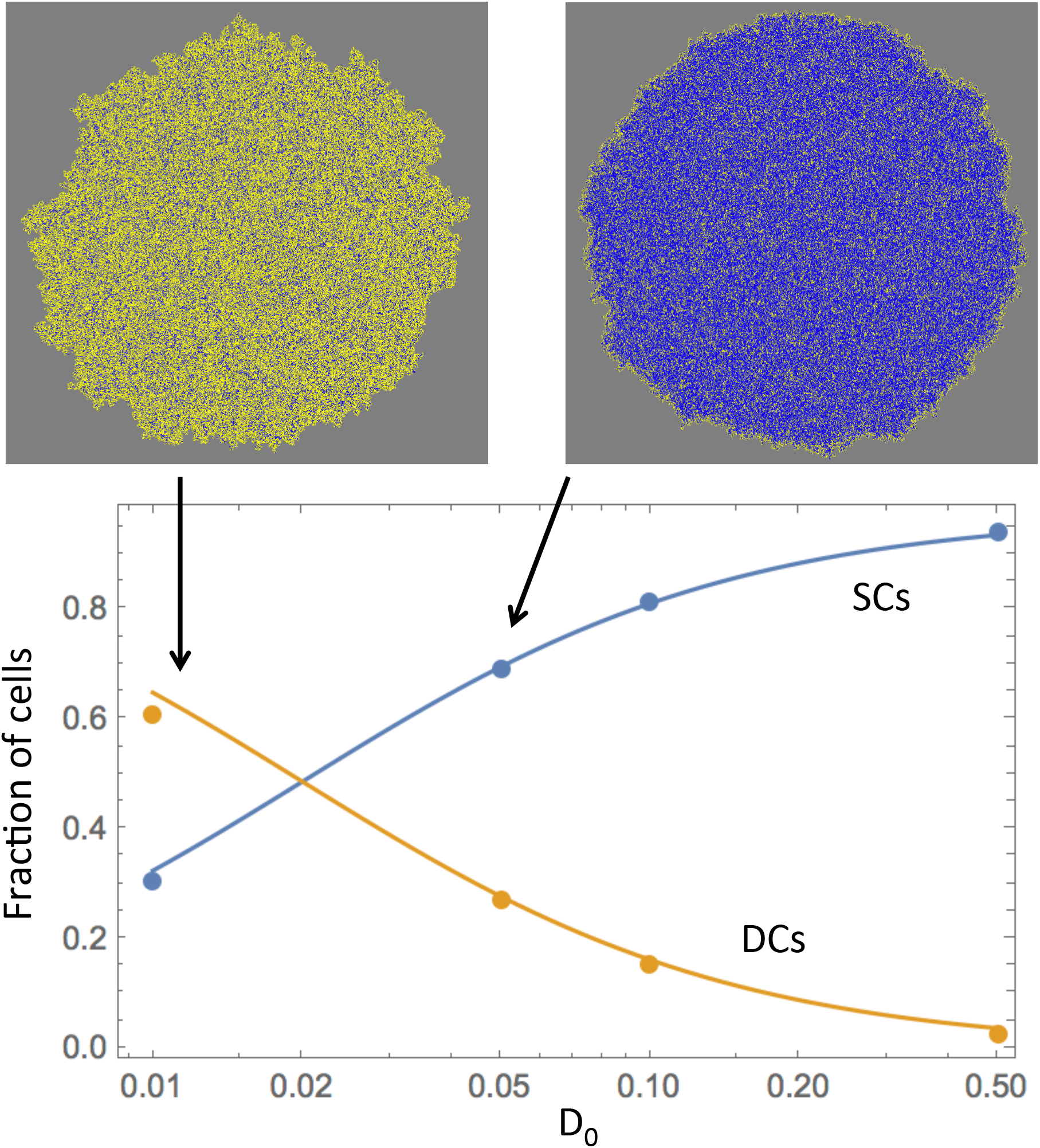
Agent based model simulations and theory. The fractions of SCs and DCs are shown for different values of parameter D_0_. Theoretical predictions from mean field modeling are given by solid lines and agent based model simulation results by points. Typical simulation results are presented for two of the parameter combinations. Gray is empty spots, blue SCs and yellow DCs. The rest of the parameters are: L_0_=0.95, P_0_=0.6, δ_0_=0.005.

Using the mean-field model, we have also calculated the degree of SC enrichment. The higher the SC death rate, the lower the overall density and the higher the fraction of DCs in the population. As time increases, the fraction of SCs reaches the value given by

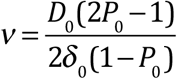

In particular, in the absence of SC death (δ_0_=0), the equilibrium value of v is infinity, which means that SCs exclude DCs everywhere in the core of the expanding range, such that DCs only concentrate in the exterior part of the colony. This corresponds to unlimited SC enrichment.

Agent based simulations were repeated assuming a 3-dimensional space, where offspring cells could be placed into one of the 27 nearest neighboring spots (Supplement). Because there are more neighbors in three dimensions, the system is more mixed, and the extent of stem cell enrichment is generally lower. (As mentioned in the beginning of this paper, no stem cell enrichment occurs in a perfectly mixed system in the absence of explicit feedback mechanisms).

## Discussion and Conclusions

The process of SC enrichment can be an important determinant of non-genetic drug resistance in cancer therapy, since CSCs are less susceptible to a variety of drugs, including chemotherapy and targeted treatment approaches [10–12]. Understanding the mechanisms that contribute to CSC enrichment is therefore a crucial step for the design of treatments that seek to sensitize tumors to drugs. SC enrichment as a cause for treatment failure has been suggested in bladder cancer [15], and might also be a reason for the occurrence of “primary resistance” of chronic myeloid leukemia (CML) to tyrosine kinase inhibitors [14].

Previous work suggested that regulatory feedback loops might contribute to CSC enrichment [13]. Feedback mechanisms have been shown to be active in tumors [8], and quantitative analysis of tumor growth data further suggests feedback regulation to influence tumor growth dynamics [16]. Here, we have provided a comprehensive analysis of how different feedback loops that might operate in tumors determine the fraction of stem cells during somatic evolution in healthy tissue or during tumor growth. Our findings indicate that the exact configuration of feedback mechanisms in tumors might be central to determining the responsiveness of tumors to therapy.

In contrast to previous approaches, we employed axiomatic modeling techniques, where model parameters such as the division rate, death rate, and self-renewal probability of cells were general functions of the number of stem and differentiated cells, which can potentially secrete feedback factors. We have thus considered a class of mathematical models that incorporate a large variety of possible feedback loops that regulate cell fate decisions in lineages. The models have the flexibility as to which cell populations exert the feedback signals, and whether feedback loops are positive or negative. The (unknown) exact details of the feedback functions remain unspecified. In the models, a mutant population is introduced which takes over by expanding from low numbers. The focus of the investigation was to establish conditions leading to stem cell enrichment in the mutant population. Some general patterns of behavior were established.

Whether or not stem cell enrichment is observed in the system depends on how the ratio D/L (deaths to divisions) changes with the population size. If it increases as the population size increases, which corresponds to either a negative feedback on L or a positive feedback on D, exerted either by stem or by differentiated cells, then SC enrichment is observed. On the other hand, if the ratio D/L decreases as the population grows, then no stem cell enrichment is predicted, and in fact SC depletion can be observed. A simple analytical approximation is derived for the long-term behavior of the expected fraction of SCs in the population.

Further, according to the model, three separate factors can decrease and even altogether prevent SC enrichment. One is the presence of SC death; the second is a very weak feedback (or no feedback) on SC divisions remaining in the mutant populations, and the third is a relatively low self-renewal probability of mutant cells.

These results generalize previous work on the determinants of stem cell enrichment, which was motivated by the experimental observation that differentiated cell death during chemotherapy of bladder cancer results in a phase of stem cell repopulation, mediated through a PGE_2_-induced would healing response [8]. Mathematical analysis suggested that in order to maintain this stem cell enrichment beyond the treatment phase, and thus to account for elevated stem cell fractions and a reduced response upon initiation of a new phase of chemotherapy, feedback mechanisms must continue to operate during untreated growth [13]. A particular negative feedback loop (from differentiated cells on the rate of stem cell division) was shown to have this effect. Here, we have shown that several different kinds of negative and positive feedback loops can have the same effect as long as the above-summarized conditions are met. This represents a first step in the quest to identify specific feedback loops in tumors that can drive CSC enrichment and hence a loss of therapy response over time. The modeling framework provided here can guide and narrow down the screening for appropriate feedback factors in specific tumors. Identification of feedback factors that are responsible for CSC enrichment is in turn important for devising treatment strategies aimed at improving the sensitivity of tumors to therapies.

The spatial models considered here have shown that in the presence of spatially restricted cell division, CSC enrichment can occur even in the absence of explicit feedback loops mediated by signaling molecules. It is the nature of spatially restricted cell spread that cells in the interim of an expanding population compete for space, such that their reproduction rate declines with population size and/or death rate increases. This has the same effect on the dynamics as the presence of explicit feedback loops. Whether this represents a physiologically important mechanism that drives CSC enrichment remains to be explored further. According to our models, stem cell enrichment is most pronounced in a 2-dimensional setting, and considerably less pronounced in a 3-dimensional setting, which applies to many solid tumors. In addition, even if spatially restricted cell divisions occur in a tumor, the presence of cell migration can destroy the reported effect (because cell migration essentially leads to a higher degree of cell mixing).

The present study provides a solid theoretical basis for implicating the presence of feedback regulatory loops as a determinant of responses to cancer therapy. This adds to the mathematical literature quantifying the role of feedback regulation for tissue and tumor dynamics [2,13,17,23,26,30–34], and builds upon the wider mathematical literature concerned with the dynamics of hierarchically structured cell populations, e.g. [35–41].

All modeling approaches contain simplifying assumptions, and the models presented here are no exception. To gain analytical insights, we reduced the complexity of the lineage differentiation pathway to include only stem cells and differentiated cells, ignoring intermediate transit amplifying cell populations with limited self-renewal capacity. Our previous work included models that explicitly took into account transit amplifying cells [13], and the relationship between the presence of negative feedback on stem cell division and the occurrence of stem cell enrichment remained qualitatively the same [42]. Other modeling approaches have treated the cell differentiation pathway as a continuous process using partial differential equations, rather than considering discrete cell sub-populations [43]. Future work is required to determine to what extent the dynamics explored here remain robust in those types of models.

## Acknowledgements

We would like to thank David Axelrod for valuable suggestions about the manuscript. We acknowledge partial funding from the National Institutes of Health (NIH) through grant 1U54CA217378-01A1 for a National Center in Cancer Systems Biology at the University of California, Irvine, and NIH grant P30CA062203 for the Chao Comprehensive Cancer Center at the University of California, Irvine. In addition, NK and DW acknowledge NIH grant 1U01CA187956, and JL acknowledges partial support from the National Science Foundation, Division of Mathematical Sciences under grant NSF-DMS-1714973.

